# Predominant tetraploidy and lack of ploidy-associated genetic structure across invasive *Lantana camara* populations in India

**DOI:** 10.64898/2026.03.11.710965

**Authors:** P. Praveen, Uma Ramakrishnan

**Affiliations:** National Centre for Biological Sciences - TIFR, GKVK Campus, Bellary Road, Bangalore 560065, India

**Keywords:** Invasive species, Polyploidy, Invasion genomics, *Lantana camara*

## Abstract

Polyploidization is widely recognised as a major driver of plant diversification, with many species persisting as mixed-ploidy systems where multiple cytotypes co-exist. Polyploids are disproportionately represented among invasive species, yet their role in facilitating biological invasions remains poorly understood. *Lantana camara*, one of the world’s most successful invasive plants, exhibits remarkable cytotype diversity, but the distribution and evolutionary relationships of these cytotypes in its native and invasive ranges have remained unclear. Here, we characterise ploidy variation and assess genetic differentiation among cytotypes in invasive *L. camara* populations across India. Flow cytometry of more than a thousand individuals reveals that tetraploids overwhelmingly dominate the invasive range, accounting for more than 95% of individuals, while triploids and hexaploids occur at much lower frequencies. Using genome-wide ddRAD-derived SNP markers from diploids, triploids, tetraploids, and hexaploids, we find no genetic differentiation among cytotypes. Instead, individuals of different ploidy levels cluster together across multiple genetic clusters, consistent with recurrent and potentially independent origins of polyploids. These patterns further suggest that *L. camara* polyploids likely arise via autopolyploid formation. Together, our results establish tetraploidy as the predominant cytotype in India’s invasive populations and reveal a lack of cytotype-specific genetic structure. These findings highlight the need to investigate the ecological advantages of tetraploids and the mechanisms that generate cytotype diversity, key steps toward understanding how polyploidy contributes to the invasive success of this globally important species.

## Introduction

Polyploidization is a major evolutionary process that has repeatedly shaped diversification across the tree of life (Landis et al., 2018; Wood et al., 2024). Although it occurs across many taxa (Booker et al., 2022; Van De Peer et al., 2017), plants are particularly remarkable for their tolerance to genome doubling; an estimated 15% of all plant species and more than one-third of angiosperms have undergone polyploidization in their evolutionary history (Landis et al., 2018; Wood et al., 2024). Polyploids are broadly classified into two types: autopolyploids and allopolyploids. Autopolyploids arise through genome duplication within a species. Whereas allopolyploids originate from hybridisation between related species, followed by genome duplication. Many polyploid species occur as a mixed ploidy systems, in which multiple cytotypes exist within a species.

Mixed-ploidy systems provide a powerful framework for understanding how polyploidization influences evolutionary and ecological dynamics in natural populations. Different cytotypes can vary in their genomic, cellular, physiological, and ecological features. One of the major ecological consequences of polyploidization is the variation in the geographic distribution of different cytotypes. In some species, cytotypes occupy distinct niches (Baniaga et al., 2020; Casazza et al., 2012; Garmendia et al., 2018), whereas in others, diploids and polyploids coexist (Chung et al., 2015; Gaynor et al., 2018; Pavel Trávníček et al., 2011). In many species, polyploids are more abundant than diploids, although the drivers of these patterns remain unclear (Pungaršek & Frajman, 2024). A commonly proposed explanation is a “tetraploid advantage” (Nagy et al., 2018), potentially arising from increased heterozygosity and heterosis, as well as neofunctionalisation or subfunctionalisation of duplicated genes (Comai, 2005). Experimental evidence supports this idea - for example, synthetic polyploids of Arabidopsis show that tetraploids outperform diploids as well as higher polyploids such as hexaploids and octaploids (Corneillie et al., 2019).

Understanding distribution patterns of polyploids also requires considering whether certain cytotypes possess intrinsic advantages that allow them to become more frequent. Polyploids often show enhanced tolerance to biotic and abiotic stress, enabling them to occupy habitats that are less suitable for diploids (Dai et al., 2020). As a result, cytotypes frequently exhibit distinct geographic or climatic distributions: polyploids are often common at higher elevations and latitudes, with temperature emerging as a strong predictor of cytotype occurrence (Rice et al., 2019; Zozomová-Lihová et al., 2015). These patterns underscore the importance of documenting the distribution of cytotypes across landscapes and identifying the ecological factors that influence these distributions. However, most evidence for such environmental associations comes from a relatively small number of temperate species, whereas mixed-ploidy systems in tropical species remain poorly studied.

Beyond their ecological consequences, polyploidization can also shape evolutionary trajectories. Polyploidisation can lead to genetic differentiation and speciation. Genome duplication followed by hybridization is an important mechanism for the formation of allopolyploids (Cao et al., 2023), enabling a relatively rapid formation of new species (Chalhoub et al., 2014; Kim et al., 2018; Levy & Feldman, 2022). However, divergence between diploids and polyploids can also accumulate gradually over time through the independent fixation of mutations in each lineage (Burns et al., 2021; Gordon et al., 2020). Gene duplication resulting from polyploidization creates opportunities for evolutionary innovation (Crow & Wagner, 2006; Glasauer & Neuhauss, 2014; Moore & Purugganan, 2003), allowing for divergence and potential evolution of new functions. Consequently, polyploid lineages may become genetically differentiated from their diploid relatives over time. However, mixed-ploidy species vary widely in this respect: some show clear genetic structuring between cytotypes (Karunarathne & Hojsgaard, 2021; Wang et al., 2021; Zozomová-Lihová et al., 2015), whereas others exhibit little differentiation, often due to recent or recurrent polyploid formation (Chumová et al., 2024; Dillenberger et al., 2018; Monnahan et al., 2019; Wang et al., 2021).

The prevalence and success of polyploids become especially apparent in biological invasions. Polyploidization and invasion success are positively correlated (Góralski et al., 2014). Polyploids are disproportionately represented among invasive species (Beest et al., 2012), suggesting that increased ploidy may facilitate colonisation and the establishment of new populations. Many invasive plants are mixed-ploidy species. Several studies show that introduced populations of invasive plants are often predominantly polyploid, even when their native ranges contain a mixture of both diploids and polyploids (Kubatova et al., 2008; Wan et al., 2020). This pattern suggests that genome duplication may enhance ecological performance and facilitate colonisation of novel environments, although polyploidy is unlikely to be the sole driver of invasion success. Despite these patterns, many invasive polyploids are poorly explored. This knowledge gap limits our ability to understand how polyploids spread, interact, and potentially contribute to invasion success. Additionally, the mechanisms that enable polyploids to outperform their diploid counterparts during invasion are still insufficiently understood, underscoring the need for comprehensive assessments of cytotype distribution in invasive plant populations.

Mixed-ploidy systems provide a powerful framework for examining the consequences of genome duplication and its potential role in invasion success. *Lantana camara* (Lantana hereafter) is a globally invasive species, where multiple ploidy levels have been documented both in its native and introduced ranges (Branda et al., 2007; Sanders, 1987), with cytotypes ranging from diploid to hexaploid. In India, individuals spanning from diploid to hexaploid levels have been reported previously (Ojha & Dayal, 1992). Despite these studies, the distribution of cytotypes in the invasive populations of Lantana remains unresolved. Resolving the spatial occurrence of various cytotypes can provide insights into their origins and evolutionary dynamics. Similarly, the extent of divergence among cytotypes has not yet been studied. Such information could help clarify whether Lantana is an auto- or allopolyploid, a distinction that is central to inferring the origins and relationships of its polyploid lineages. To address these gaps, we study the invasive Lantana populations in India and ask: (i) What is the predominant cytotype of Lantana in wild populations? and (ii) Do different cytotypes exhibit genetic differentiation? Answering these questions will shed light on the origins, spatial structure, and evolutionary dynamics of polyploidy in this globally invasive species.

## Materials and Methods

### Sampling

Lantana saplings were collected from six locations across India for the ploidy estimations (Supplementary Figure 1). The locations were selected from regions where Lantana was reported before 1900 to include locations of early introduction (Kannan et al., 2013). These locations are Pachmarhi, Kolkata, Pune, and areas near Ooty, such as Gudalur, Bandipur Tiger Reserve and Sathyamangalam Tiger Reserve. Within each location, saplings were collected opportunistically, mainly from roadsides, abandoned fields, and other disturbed habitats. The collected saplings were planted in the greenhouse at the National Centre for Biological Sciences, Bangalore and young leaves were collected for the ploidy estimation. A total of 1070 plants were tested for their ploidy. For genomic analysis, 46 samples were used from five locations (Sathyamangalam Tiger Reserve, Gudalur, Bandipur Tiger Reserve, Jodhpur, and Jaipur) with varying cytotypes, including those previously sampled for ploidy estimation. Fresh leaf tissue from these samples was used for DNA extractions. Some samples from western India that were not included in the ploidy survey were also incorporated into the genomic dataset.

### Ploidy estimation

Plant ploidy levels were estimated following the protocol described by Dolezelt et al. 1998. Briefly, young rapidly growing leaves were used to minimise the interference from phenolic compounds on ploidy estimation. Approximately 20mg of leaf tissue was chopped in 500 µl of ice-cold Otto 1 solution using a sharp scalpel. The homogenate was mixed and filtered through a 40-micron nylon mesh. The filtrate was centrifuged at 150g for five minutes. After discarding the supernatant, the nuclei pellet was resuspended with 50 µl fresh Otto 1 solution and 500 µl of Otto 2 buffer was added to the mixture. RNase and Propidium iodide solutions were added at the final concentration of 50 and 25 µg/ml, respectively. The samples were incubated for five minutes, and the fluorescence emitted from the nuclei was recorded using BD LSRFortessa and BD FACSvers flow cytometers. Chicken Erythrocyte Nucleus (CEN) was used as a standard to calculate the DNA content. Based on previous reports, tetraploid Lantana individuals contain approximately 3 pg of nuclear DNA (Deng & Wilson, 2017; Deng et al., 2017),and this value was used as the reference standard for ploidy determination.

### DNA extractions and ddRAD library preparation

Fresh leaf samples were collected from plants, freeze-dried using liquid nitrogen, and ground into fine powder using a mortar and pestle. DNA was extracted using Qiagen DNeasy Plant Kit using the manufacturer’s instructions. The concentration of the DNA was measured using a Qubit fluorometer, and the DNA was stored at -20^0^C until it was used for sequencing. ddRAD libraries for the extracted samples were prepared following (Tyagi et al., 2024). Briefly, restriction digestion was performed using SphI and MlucI enzymes. Adapters specific to the restriction sites were ligated, followed by the ligation of individual-specific indexes. Libraries were size-selected using magnetic beads to isolate fragments ranging from 200 to 500 bp. Finally, all the individual libraries were pooled to create a 2 nM final library, which was sequenced on either the NovaSeq6000 or HiSeq2500 platform.

### Genomic analysis

The raw reads were demultiplexed, and the adapter sequences and low-quality reads were removed using Trimmonatic - v.0.33.0 (Bolger et al., 2014). The trimmed reads were then aligned to the Lantana reference genome (Adwait G. Joshi, 2022) using BWA-MEM (Li & Durbin, 2009). The variant calling was performed with Freebayes (Garrison & Marth, 2012), specifying the ploidy of each individual. The resulting vcf file was filtered using Samtools (Li, 2011), applying various filtering criteria. The filtered vcf was used for downstream analysis of diversity parameters. PLINK (Purcell et al., 2007) was used to generate eigenvalue and eigenvector files for the PCA. For constructing the phylogenetic network, StAMPP package was used (Pembleton et al., 2013), and the resulting tree was visualized using SplitsTree4 (Huson, 1997). F_ST_ was calculated using Genodive (Meirmans, 2020). Nei’s genetic distances between individuals were calculated using StAMPP package. We used ADMIXTURE (Alexander et al., 2009) to infer population structure. Although ADMIXTURE is not specifically designed for polyploid datasets, the results obtained were consistent with patterns recovered from other population-differentiation analyses. We subsampled alleles at each site to generate diploid-like genotypes using a custom script, following approaches implemented in previous studies. The Admixture results were visualised using CLUMPAK (Kopelman et al., 2015) and the optimum K was selected based on Evanno’s delta K method (Evanno et al., 2005).

## Results

### Tetraploidy is the predominant cytotype in India’s invasive Lantana camara populations

Plant samples from six locations were screened for their ploidy using flow cytometry (Supplementary Figure 1). Chicken erythrocyte nuclei were used as a standard to measure the genomic DNA content. Based on this standard, the DNA content of a reference plant from Bangalore - a wild-grown tetraploid was measured (with 3pg nuclear DNA content) and used as the control for further analysis. Ploidy levels of all the other samples were determined relative to this control. Four ploidy levels from diploid to hexaploid were detected in the wild populations (Figure 1A). Across the 1,070 plants analysed, the vast majority (1,022; 95.5%) were tetraploid (Figure 1B, Supplementary Table 1). Other cytotypes were also detected, including 18 triploids and 29 hexaploids (Figure 1B). Notably, some locations, such as Gudalur and Sathyamangalam, exhibited a relatively high proportion of non-tetraploid cytotypes (Figure 1C). In Gudalur, 6.46% of the plants were triploids, and 6.89% were hexaploids, while in Sathyamangalam, 4.86% were hexaploids (Figure 1C, Supplementary Table 1). Two diploid individuals were detected in western India; however, these were only included in the genomic analysis.

**Figure 1.**
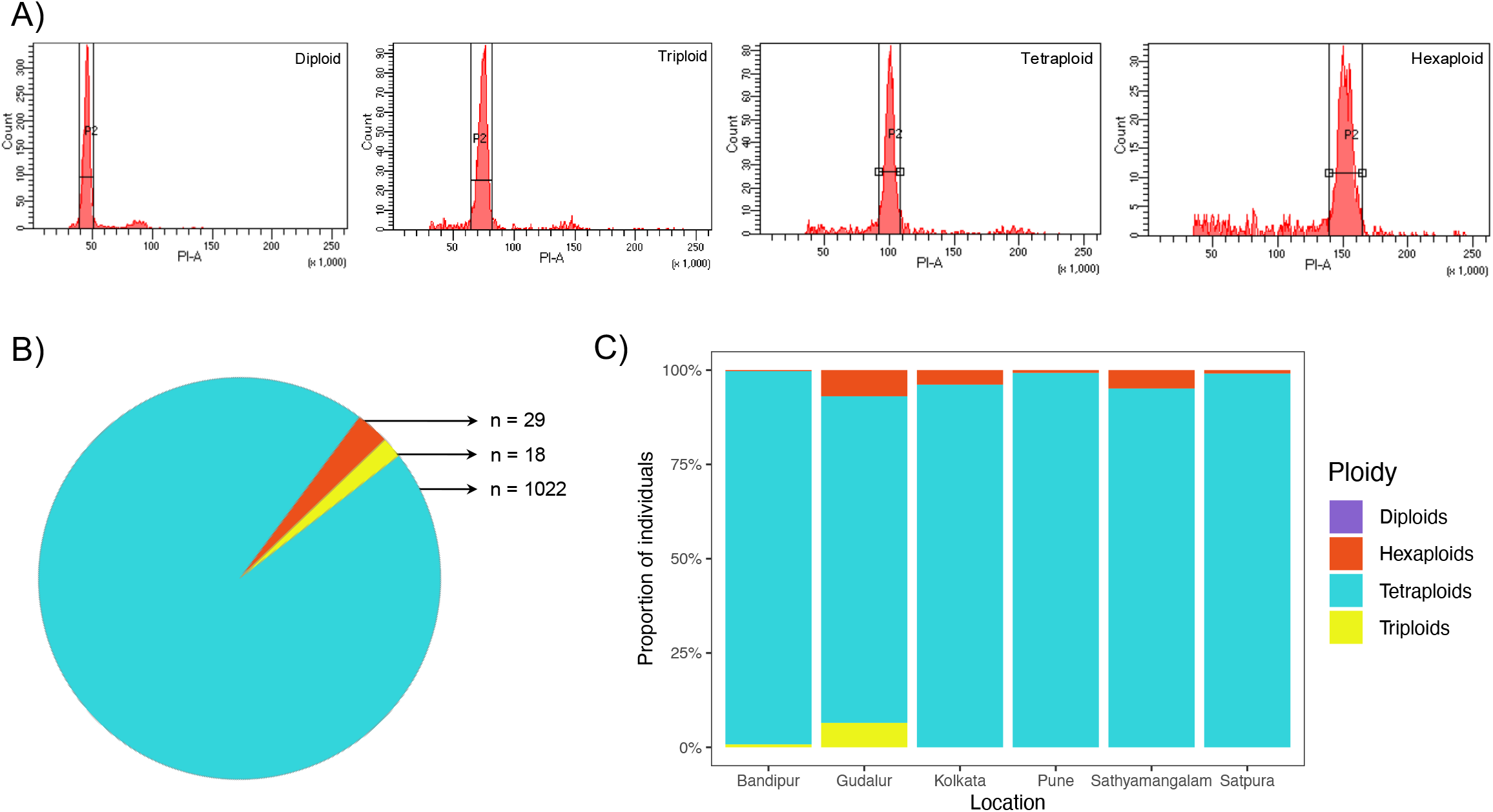
Cytotype diversity of invasive Lantana camara. A) Representative flow cytometry histograms showing fluorescence peaks for different cytotypes. B) Proportion of each cytotype across all sampled individuals C) Site-wise distribution of cytotypes showing variation across locations

### No genetic differentiation among Lantana cytotypes, despite the presence of multiple genetic clusters

To assess genetic differentiation among cytotypes, we sequenced 46 individuals using ddRAD-seq, including two diploid, nine triploid, 18 tetraploid and 17 hexaploid plants. Genetic differentiation among cytotypes was examined using PCA, *Admixture*, F_ST_ and phylogenetic networks. In the PCA, the first and second principal components explained 22.1% and 18% of the genetic variation, respectively (Figure 2A). Notably, individuals did not cluster according to ploidy. Instead, multiple genetic clusters were detected within cytotypes, and several clusters contained individuals of different ploidy levels. For example, some clusters included triploid, tetraploid and hexaploid individuals. The two diploid individuals from western India clustered with the tetraploid individuals from the same region, further supporting the absence of ploidy-linked genetic differentiation. The PCA using other PC combinations (PC1-PC3 and PC2-PC3) also showed similar trends (Supplementary Figure 5).

**Figure 2.**
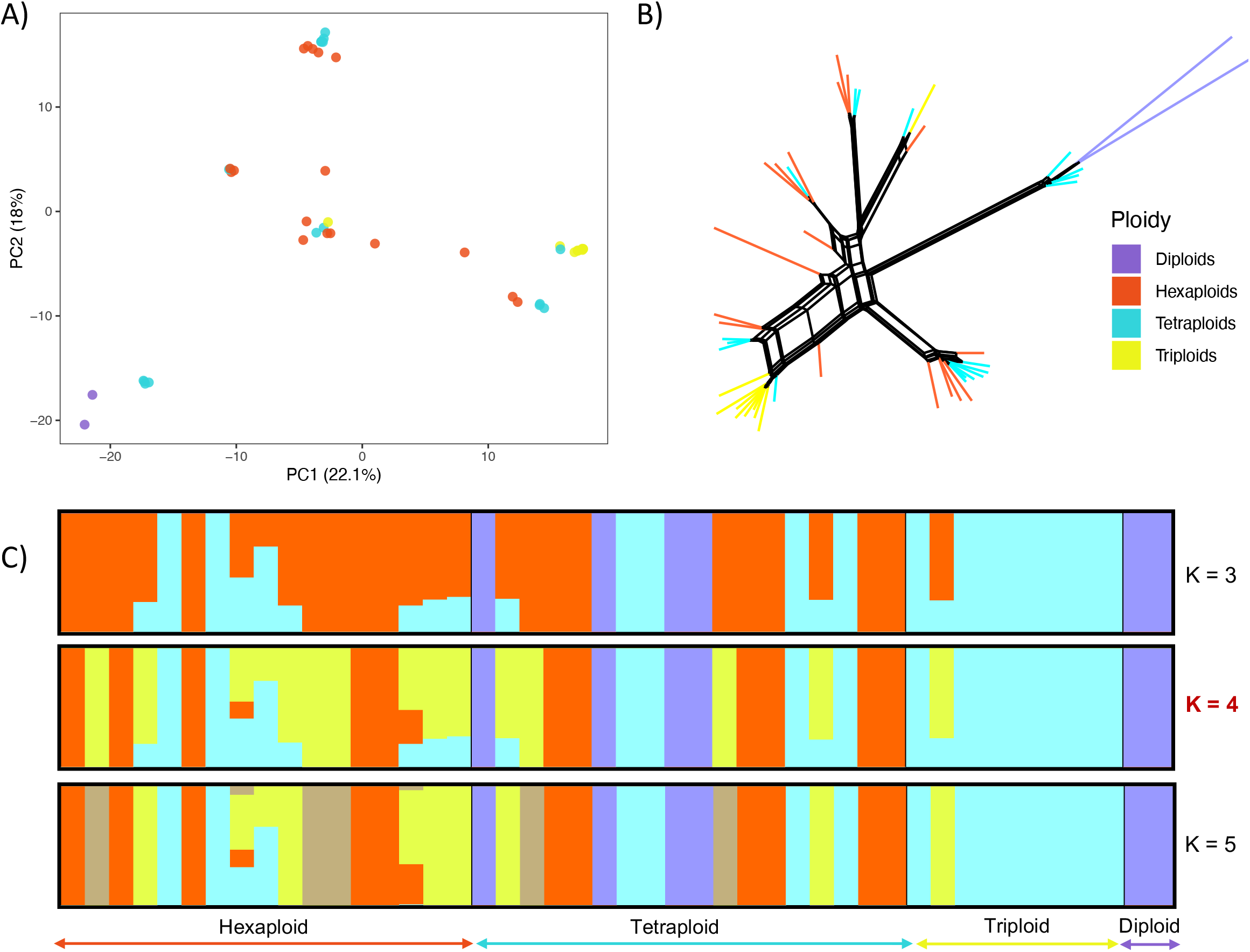
Patterns of genetic differentiation among invasive Lantana camara cytotypes A) Principal component analysis (PCA) showing genetic clustering among cytotypes B) Phylogenetic network C) Ancestry proportions inferred using ADMIXTURE

In the phylogenetic network, a similar pattern was observed: individuals did not form clades based on ploidy (Figure 2B). Instead, monophyletic groups often included individuals from different cytotypes, further indicating a lack of genetic separation among them. Although diploid individuals exhibited longer branch lengths - suggestive of greater genetic divergence - they nonetheless clustered with tetraploids from the same geographic region. Pairwise *F*_ST_ values corroborated these patterns, showing very weak differentiation among cytotypes, with the highest *F*_ST_ (0.047) observed between triploids and hexaploids (Supplementary Table 2). The heatmap showed a similar pattern, with the diploids being the most genetically distinct (Supplementary Figure 4).

The population structure of Lantana inferred using Admixture showed no clustering pattern associated with cytotype. The Evanno’s ΔK method indicated that K = 4 was the most likely number of genetic clusters (Supplementary Fig. 2). Individuals from the same ploidy level were assigned to multiple clusters, and conversely, each cluster contained individuals belonging to different cytotypes. A similar lack of structure was observed geographically, with individuals belonging to different genetic clusters broadly distributed across sampling locations rather than being region-specific (Supplementary figure 3).

## Discussion

Polyploidization is widely regarded as an evolutionary strategy that aided the diversification of many lineages (Han et al., 2020), yet polyploid systems, especially in the tropics, remain comparatively understudied. Invasive polyploids are particularly valuable to study, as they can shed light on the ecological advantages associated with genome duplication. In this study, we examined the invasive *Lantana camara* populations across India to characterise cytotype distribution in the wild and assess genetic differentiation among cytotypes. We detected the presence of various cytotypes in the wild populations, with tetraploids being the most abundant cytotype. Genomic analysis revealed a lack of clear genetic differentiation among cytotypes, alongside the presence of multiple genetic clusters within certain cytotypes. Together, these patterns suggest multiple, potentially independent, autopolyploid origins in Lantana. Genomic analyses have historically been limited by the complexity of polyploid genomes; recent methodological advances are helping to overcome these limitations, and this study also tried to utilise this advancement.

### Prevalence of tetraploidy in Lantana populations

Many plant species exhibit variation in ploidy level, and Lantana is a notable example of such a mixed-ploidy system. Our findings reveal that tetraploids dominate the invasive populations in India. Although polyploidy has been documented previously in Lantana - both in wild plants and horticultural cultivars (Czarnecki et al., 2014; Ojha & Dayal, 1992; Spies, 1984) - comprehensive surveys across its native and introduced ranges remain scarce. Our study provides the first broad-scale evidence that tetraploids dominate invasive populations in India. However, the global distribution of cytotypes remains incomplete due to the lack of comprehensive ploidy estimates. Tetraploids in other species have been reported to exhibit an advantage over other cytotypes, although the exact reason for such a pattern remains unknown (Nagy et al., 2018). For example, synthetic tetraploids of *Arabidopsis thaliana* have demonstrated increased productivity (Corneille et al., 2019). While such factors may contribute to the dominance of tetraploids in Lantana, the underlying mechanisms require targeted experimental validation.

Polyploids are disproportionately represented among invasive species (Beest et al., 2012; Pandit et al., 2011), and our results show that Lantana follows this broader pattern. Polyploidy is known to enhance ecological tolerance and facilitate colonisation of novel environments (van de Peer et al., 2021), and similar advantages may contribute to the success of tetraploid Lantana during its spread. Alternatively, the high prevalence of tetraploids in India could reflect introduction bias, where particular cytotypes were introduced more frequently or naturally selected in the introduced range. Distinguishing between these possibilities will require a systematic assessment of cytotype distributions in Lantana’s native range.

### Cytotype diversity in Lantana populations

Indian Lantana populations exhibit cytotype diversity, with individuals of diploid, triploid, and hexaploid cytotypes occurring alongside the predominant tetraploid cytotype. However, the factors driving the production of these diverse cytotypes remain unclear, but the rarity of non-tetraploid cytotypes suggests that they may arise from tetraploid lineages through processes such as unreduced gamete formation. The absence of strong genetic differentiation among cytotypes in our dataset is consistent with this scenario, indicating recent or recurrent polyploid formation. In Lantana, the production of unreduced gametes has been linked to the formation of polyploids (Ii & Deng, 2009). Also, an increase in unreduced gamete production is known to lead to greater cytotype diversity in other plant systems (Kreiner et al., 2017; Mason & Pires, 2015). This evidence further supports our speculation, although further validation is required.

The presence of non-tetraploid cytotypes in regions such as Gudalur and Sathyamangalam in the Western Ghats is particularly intriguing. These areas are cooler and wetter, raising the possibility of the potential influence of environmental factors on the generation of polyploids in Lantana. Previous research has shown that environmental factors can significantly impact the production of unreduced gametes (Kreiner et al., 2017). In species like *Cardamine amara*, climatic factors such as temperature, photosynthetically active radiation and annual precipitation have been shown to influence the occurrence of tetraploids (Zozomová-Lihová et al., 2015). Assessing unreduced gamete production and cytotype frequencies across environmental gradients in the Western Ghats and surrounding regions would therefore provide valuable insight into whether ecological factors contribute to polyploid formation in Lantana.

### Genetic differentiation among cytotypes of Lantana

Understanding the genetic differentiation among different cytotypes can provide insights into their origin and evolutionary trajectories. In many plant species, polyploidization is accompanied by genetic divergence and can act as a pathway to speciation (Cao et al., 2023). Our genomic analyses of Lantana reveal no detectable genetic differentiation among cytotypes. This pattern may reflect the recent origin of these cytotypes, combined with continued gene flow among them. Despite the absence of cytotype-level differentiation, we detected multiple genetic clusters that each contain mixtures of triploids, tetraploids, and hexaploids. This structure suggests that polyploid cytotypes may have arisen repeatedly and independently from genetically distinct lineages - an interpretation consistent with recurrent autopolyploidization through the fusion of unreduced gametes (Praveen et al, 2025). The production of unreduced gametes has been documented in Lantana (Ii & Deng, 2009), and repeated formation of autopolyploids is well known in other species (Monnahan et al., 2019). Further, previous work indicates that Lantana can undergo self-fertilisation, and the different genetic lineages can be inbred lines formed through this process (Praveen et al., 2025). Over longer evolutionary timescales, autopolyploids may accumulate mutations leading to divergence, but our results suggest that such differentiation has not yet occurred in Lantana cytotypes.

While the present study provides robust insights into ploidy distribution and genetic differentiation, a few aspects warrant consideration. Direct chromosome counts of the flow cytometry standards would have provided an additional layer of validation, but we relied on previous nuclear DNA content estimates. In addition, inclusion of the native range population would enable stronger inferences about invasion-associated change in cytotype composition, an important direction for future research. Finally, more uniform sampling across ploidy levels in the genomic analysis would further strengthen comparisons among cytotypes.

## Conclusions

Together, this study provides the first comprehensive assessment of cytotype distribution and genetic differentiation among cytotypes in invasive Lantana populations. We show that tetraploids overwhelmingly dominate wild populations, while other cytotypes occur at low frequencies. We detect no genetic differentiation among different cytotypes; instead, multiple mixed-cytotype genetic clusters point to the chances of autopolyploidization, likely driven by the fusion of unreduced gametes. These patterns suggest that polyploidization in Lantana is ongoing and not yet accompanied by substantial genomic divergence. By documenting both the prevalence of polyploidy and the lack of cytotype-level structure in this invasive tropical species, our study highlights the need for expanded sampling across the native range and other invaded ranges. Our findings provide fresh insight into a potential mechanism linking polyploidization to invasion success and open new avenues to explore how these processes interact in Lantana.

## Acknowledgements

We are thankful to Keval Palya, Mamta M., Mayuresh Gangal, Kannan and Rajat Rastogi for their help with sampling and laboratory experiments. We thank the forest departments of Karnataka (No: PCCF (WL) /E2/CR-52/2019-20), Tamil Nadu (No: WL(A)/52852/2019) and Madhya Pradesh (MP permit no:835, dated: 30-01-2020) for providing necessary permits. PP was supported by NCBS/TIFR (Department of Atomic Energy). This work was supported under DBT project No-BT/PR29251/FCB/125/18/2018 and the NCBS-TIFR internal plan fund that was awarded to UR. The NCBS data cluster used is supported under project 12-R&D-TFR-5.04-0900, Department of Atomic Energy, Government of India. For institutional support, we thank the National Centre for Biological Sciences (NCBS). We thank the Next Generation Genomics facility at NCBS, for helping with the sequencing.

## Author Contributions

Conceptualisation: P.P., and U. R., Data collection: P.P., Laboratory work: P. P., Data analysis: P.P, Project administration: P.P., and U.R., Funding acquisition: U. R, Writing original draft: P.P., Writing – Review and editing: U.R., Supervision: U.R.

## Data availability statement

Accession numbers for the raw ddRAD sequencing data and the GitHub link for analysis codes will be provided upon acceptance of the manuscript.

