## Supplementary information for "Predominant tetraploidy and lack of ploidy-associated genetic structure across invasive *Lantana camara* populations in India"

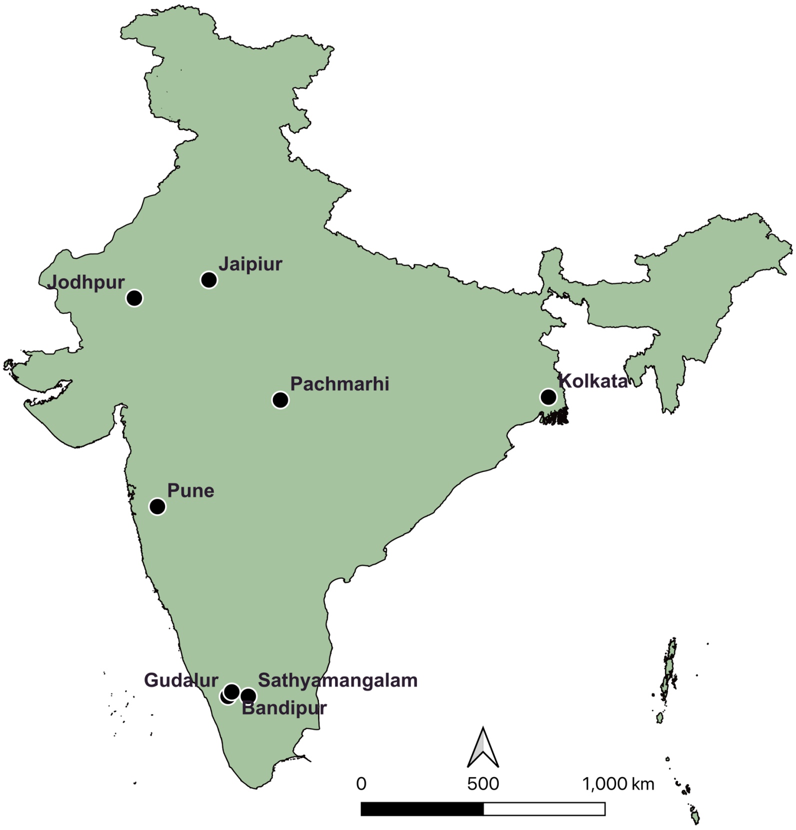


*Figure 1 Geographic locations of* Lantana *populations sampled for the study*

*Table 1 Distribution of cytotypes recorded at each location*

| | Location | Diploids | Triploids | Tetraploids | Hexaploids | Total |
|---|---|---|---|---|---|---|
| 1 | Sathyamangalam TR | - | - | 176 | 9 | 185 |
| 2 | Bandipur TR | - | 3 | 375 | 1 | 379 |
| 3 | Gudalur | - | 15 | 200 | 16 | 232 |
| 4 | Pune | - | - | 136 | 1 | 137 |
| 5 | Satpura | - | - | 110 | 1 | 111 |
| 6 | Kolkata | - | - | 25 | 1 | 26 |
| | **Total** | | 18 | 1022 | 29 | **1070** |

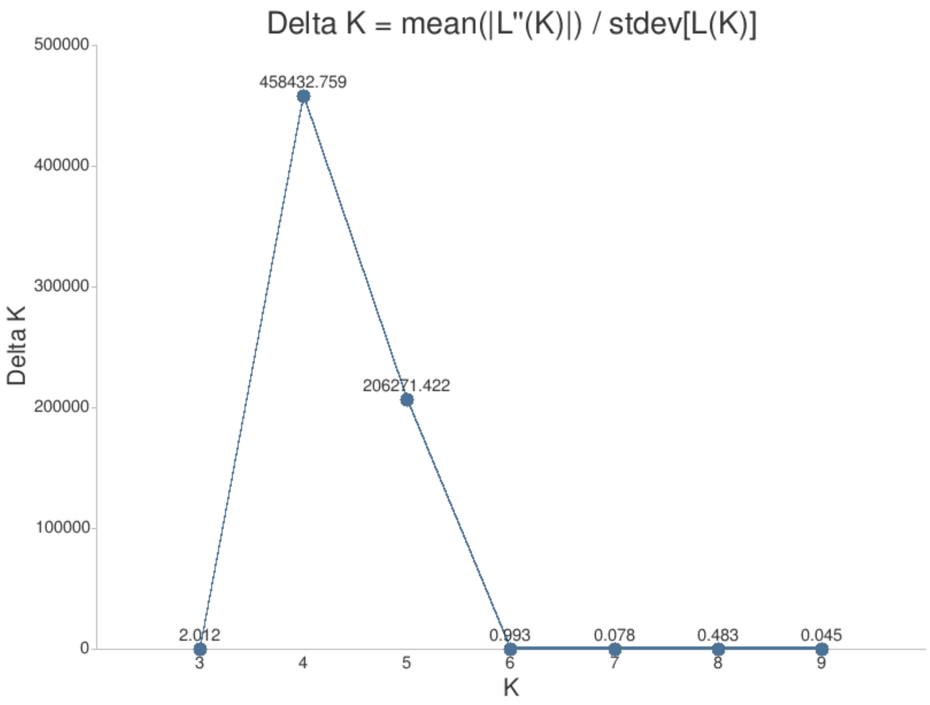


*Figure 2 Optimal K inferred using Evanno's ΔK method. The K value with the highest ΔK is considered the best-supported number of genetic clusters*

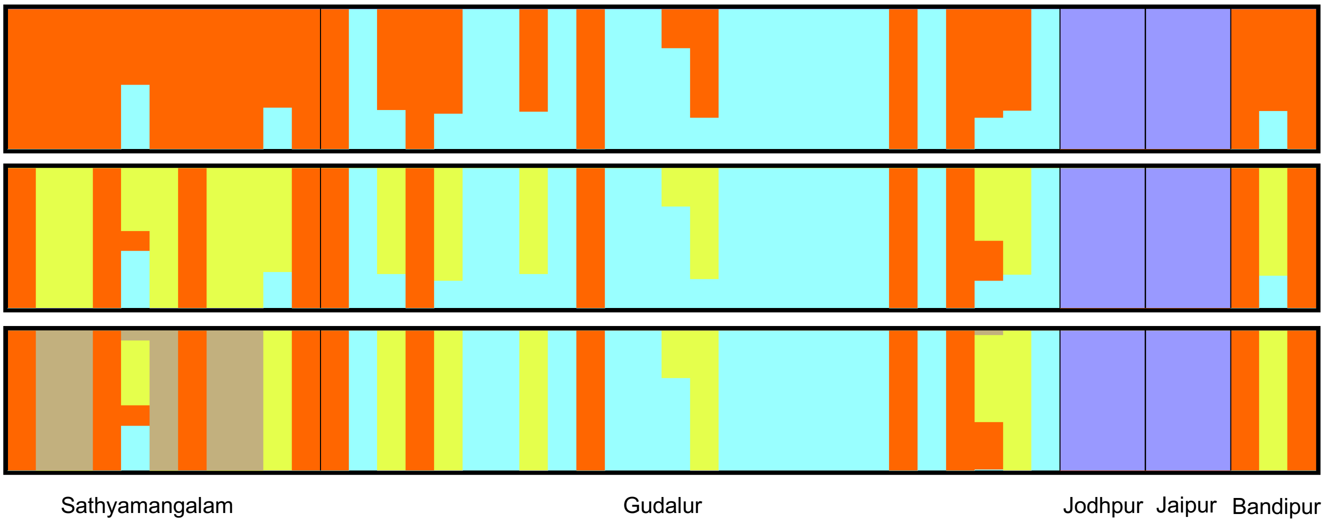


*Figure 3 Population structure of* Lantana *cytotypes, arranged by population of origin*

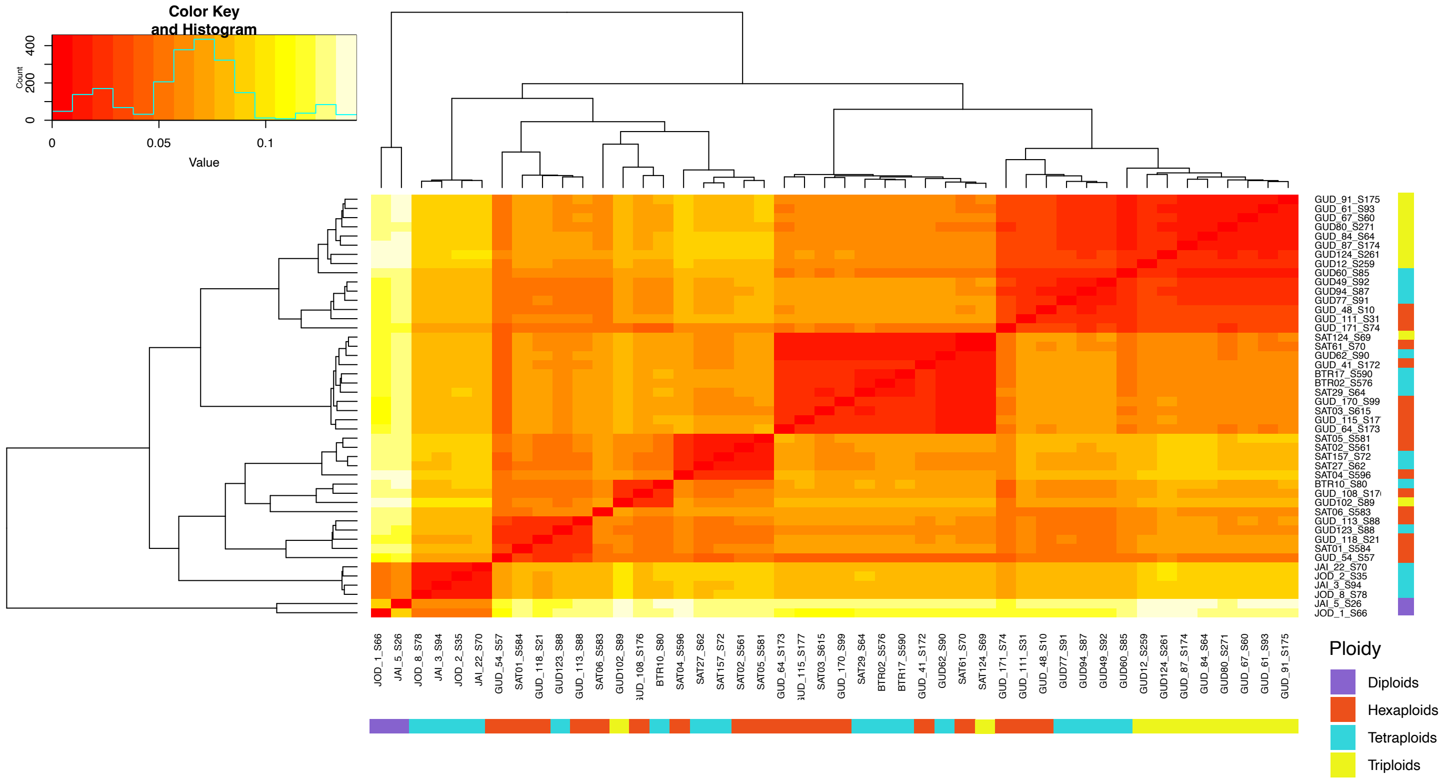


*Figure 4 Heatmap of Nei's genetic distances among all* Lantana *samples. The colour bar denotes the ploidy level of each individual*

*Table 2 Pairwise FST values between different cytotypes of* Lantana

| | **Diploid** | **Triploid** | **Tetraploid** | **Hexaploid** |
|---|---|---|---|---|
| **Diploid** | 0 | 0 | 0 | 0 |
| **Triploid** | | 0 | 0.041 | 0.047 |
| **Tetraploid** | | | 0 | 0.003 |
| **Hexaploid** | | | | 0 |

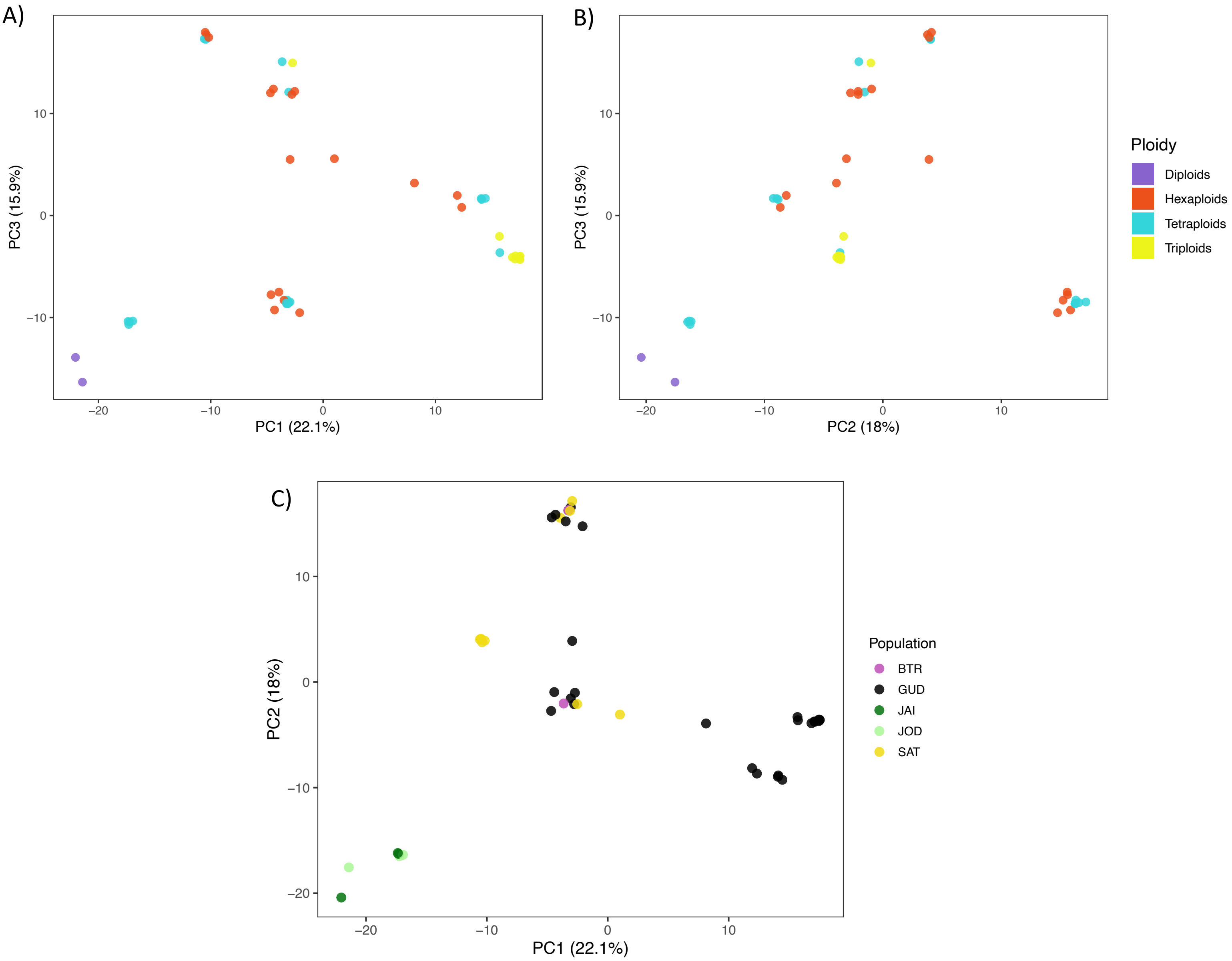


*Figure 5 Principal component analysis (PCA) illustrating genetic clustering among different cytotypes, based on PC1–PC3 (A) and PC2–PC3 (B) axes. PCA showing genetic clustering of different cytotypes, with samples coloured according to their population of origin (C)*
